# Glucose PTS Modulates Pyruvate Metabolism, Bacterial Fitness, and Microbial Ecology in Oral Streptococci

**DOI:** 10.1101/2022.09.21.508965

**Authors:** Lin Zeng, Alejandro R. Walker, Robert A. Burne, Zachary A. Taylor

**Author notes:** Corresponding author. University of Florida, Department of Oral Biology, 1395 Center Dr., Gainesville, FL 32610.

## Abstract

Spontaneous mutants with defects in the primary glucose phosphotransferase (PTS) permease (*manLMNO*) of *Streptococcus sanguinis* SK36 showed enhanced fitness at low pH. Transcriptomics and metabolomics with a *manL* deletion mutant (SK36/*manL*) revealed redirection of pyruvate to production of acetate and formate, rather than lactate. The observations were consistent with measurements of decreased lactic acid accumulation and increased excretion of pyruvate and H_2_O_2_. Genes showing increased expression in SK36/*manL* included those encoding carbohydrate transporters, extracellular glycosidases, intracellular polysaccharide (IPS) metabolism, arginine deiminase, and pathways for metabolism of acetoin, ethanolamine, ascorbate and formate; along with genes required for membrane biosynthesis and adhesion. *Streptococcus mutans* UA159 persisted much better in biofilm co-cultures with SK36/*manL* than with SK36, an effect that was further enhanced by culturing the biofilms anaerobically but dampened by adding arginine to the medium. We posited that the enhanced persistence of *S. mutans* with SK36/*manL* was in part due to excess excretion of pyruvate by the latter, as addition of pyruvate to *S. mutans-S. sanguinis* co-cultures increased the proportions of UA159 in the biofilms. Reduction of the buffer capacity or increasing the concentration of glucose benefited UA159 when co-cultured with SK36, but not with SK36/*manL*; likely due to the altered metabolism and enhanced acid tolerance of the mutant. When *manL* was deleted in *S. mutans* or *Streptococcus gordonii,* the mutants presented altered fitness characteristics. Our study demonstrated that PTS-dependent modulation of central metabolism can profoundly affect streptococcal fitness and metabolic interactions, revealing another dimension in commensal-pathogen relationships influencing dental caries development.

**Importance:** Dental caries is underpinned by a dysbiotic microbiome and increased acid production. As beneficial bacteria that can antagonize oral pathobionts, oral streptococci such as *S. sanguinis* and *S. gordonii* can ferment many carbohydrates, despite their relative sensitivity to low pH. We characterized the molecular basis for why mutants of glucose transporter ManLMNO of *S. sanguinis* showed enhanced production of hydrogen peroxide and ammonia, and improved persistence under acidic conditions. Significant metabolic shift involving more than 300 genes required for carbohydrate transport, energy production, and envelope biogenesis was observed. Significantly, *manL* mutants engineered in three different oral streptococci displayed altered capacities for acid production and interspecies antagonism, highlighting the potential for targeting the glucose-PTS to modulate the pathogenicity of oral biofilms.

## Introduction

Dental caries is an infectious disease underpinned by a dysbiotic microbiome, dental biofilms that become dominated by acidogenic and acid-tolerant (aciduric) pathobionts, including *Streptococcus mutans, Lactobacillus, Bifidobacteria, Scardovia,* and *Candida spp* (1–3). Concurrently, there is a reduction in biodiversity of cariogenic microbiomes and, in particular, in the abundance of certain commensal species, such as the mitis group of streptococci that include multiple species with strong associations with oral health (4, 5). In fact, there often exists an inverse correlation in the abundance of the aforementioned cariogenic species and that of two important commensals, *Streptococcus sanguinis* and *Streptococcus gordonii;* with these two commensals frequently associated with dental health (6–9). Based on these studies, probiotic strategies are being explored to boost the abundance of these beneficial bacteria in the oral microbiome as a means of preventing caries (10). Considering the complexity of oral microbial communities, however, a comprehensive understanding of the molecular mechanisms responsible for shaping the dynamics between pathobionts and commensals is a prerequisite to the success of therapies that seek to modify the composition of the oral microbiome (11).

Most oral streptococci have an incomplete tricarboxylic acid (TCA) cycle and lack the ability for oxidative respiration, which requires that they use carbohydrate fermentation for energy production, an activity that often results in considerable acidification of the environment. The organic acid that is the dominant product when fermentable carbohydrates are abundant, lactic acid, is created by reduction of pyruvate by lactate dehydrogenase (Ldh)(12). Many species of mitis group streptococci, including *S. sanguinis* and *S. gordonii*, produce hydrogen peroxide (H_2_O_2_) as a function of pyruvate oxidation, and are capable of alkalinization of their cytoplasm and environment through ammonia generation by the arginine deiminase (AD) system (13). As a major caries pathogen, *S. mutans* specializes in carbohydrate fermentation under low-pH conditions, but is more sensitive to H_2_O_2_ than most commensal streptococci. While much has been done to understand the aciduricity of *S. mutans* and the peroxigenic activity of commensals, less is understood regarding adaptation to acidic environments by commensals, many of which are significant contributors of organic acids to dental biofilms, even during caries development (14, 15). There is also enormous genomic and phenotypic heterogeneity in the acidogenicity, acid tolerance, and AD expression among isolates of oral streptococci (15–17). Yet, much remains to be learned regarding the molecular mechanisms used by commensal streptococci to adapt and diversify in a microbial landscape being invaded or dominated by highly cariogenic bacteria.

Prioritization of metabolism of different carbohydrates by streptococci is controlled mainly by carbon catabolite repression (CCR), involving transcriptional regulation by catabolite control protein CcpA and/or modulation of gene expression and carbohydrate uptake by components of the sugar: phosphotransferase system (PTS)(18, 19). Notably, CcpA regulates expression of functions described above, in particular acid production in *S. mutans* (20), H_2_O_2_ production by *S. sanguinis* and *S. gordonii* (21), and AD gene expression (22). CcpA plays major roles in streptococci in the regulation of central carbon metabolism and virulence expression (20, 23–25), but in many streptococci, including *S. mutans, S. gordonii,* group A streptococci (GAS), and *Streptococcus pneumoniae*, the glucose-PTS can be a dominant regulator governing the utilization of non-preferred carbohydrates by acting independently or in concert with CcpA (26–31).

We recently identified in subpopulations of two separate stocks of *S. sanguinis* SK36 mutations of the gene encoding the EIIAB (ManL) components of the primary glucose-PTS permease (ManLMNO) that resulted in truncations of the ManL protein (32). Despite having a defect in carbohydrate transport, these mutants showed enhanced capacity in persisting and fermenting sugars at lower pH values than the parental strain. The mutants also had increased production of H_2_O_2_, which conferred a greater capacity to inhibit the growth of *S. mutans* (32). To understand the molecular basis for these observations, we conducted transcriptomic and metabolomic analyses of a *manL* mutant of SK36. When the effects of *manL* mutation were analyzed in *S. gordonii* and *S. mutans*, we observed significant changes in fitness and interbacterial competition that revealed a novel role for the glucose-PTS in regulating energy metabolism and oral microbial ecology.

## Results and Discussion

### Effects of ManL on the transcriptome of *S. sanguinis*

To begin to dissect the molecular mechanisms underlying improved acid resistance and H_2_O_2_ production by *manL* mutants (32), we performed an RNA-Seq on a strain carrying a targeted deletion of *manL* with the wild-type SK36 in cells grown on TY-glucose. At a cutoff of ≤0.01 as the *P* value and ≥2-fold change in mRNA abundance, a total of 311 genes, approximately 14% of the genome (33), were deemed differentially expressed (DE; see Fig. 1, Fig. S1, and Table S1 for details), 64% of which showed increased expression. The most prominent functions affected by the deletion of *manL* were energy metabolism and carbohydrate transport and catabolism, consistent with the reported fitness phenotype.

**Fig. 1.**
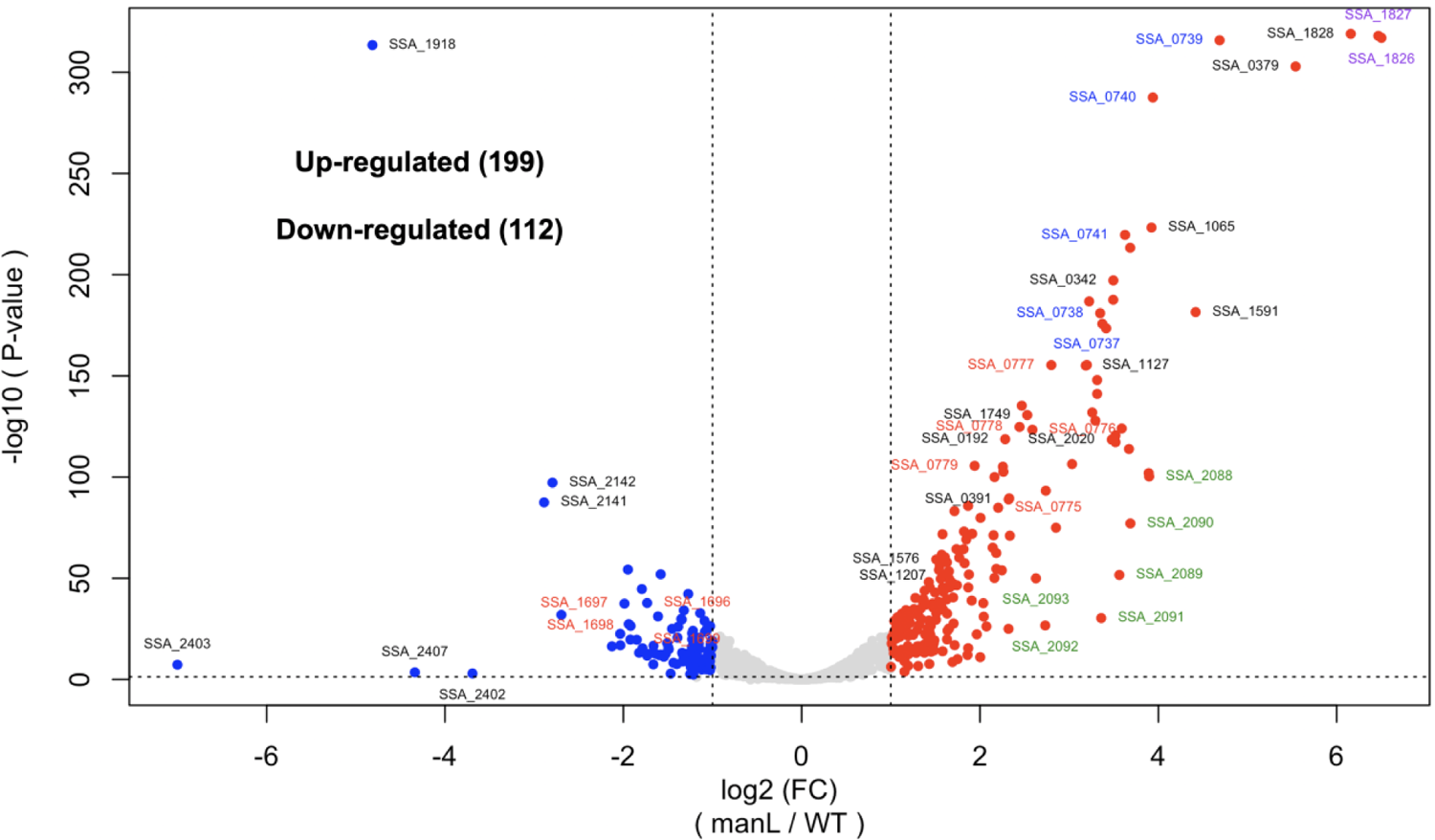
RNA-Seq: volcano plot visualizing differentially expressed genes in SK36/*manL* versus SK36. The plots represent 3 independent replicates cultured in TY-glucose to mid-exponential phase (OD_600_ = 0.5∼0.6) before being harvested for RNA extraction and deep sequencing. Genes or ORFs shown here met the following criteria: |Log_2_(*manL*/WT)| ≥1, and *P* value ≤0.01.

#### i) Central metabolism

Deletion of *manL* in SK36 altered expression of genes that were known or predicted to participate in metabolism of pyruvate or metabolites derived from pyruvate (Fig. 2). Specifically, the gene responsible for oxidizing pyruvate to create H_2_O_2_ with concomitant acetyl-phosphate (Ac-P) production, *spxB* (SSA_0391)(34), was upregulated 5-fold, whereas the pyruvate-reducing lactate dehydrogenase (*ldh,* SSA_1221) was downregulated by ∼40%. Pyruvate-formate lyase (*pfl,* SSA_0342) and a related pyruvate formate lyase-activating protein (*pflA,* SSA_1749) were upregulated 11- and 6-fold, respectively, and the pyruvate dehydrogenase operon (SSA_1137 to SSA_1140) was upregulated 2- to 3-fold. These changes in gene regulation were consistent with increased H_2_O_2_ excretion and reduced lactate production by the *manL* mutant (32), indicative of a shift from reductive to oxidative metabolism of pyruvate and production of alternative metabolites, including Ac-P, acetyl-CoA, formate, and acetate. As an important reaction to balance the intracellular NADH:NAD^+^ ratio by oxidizing NADH, the activity of Ldh is allosterically modulated by glycolytic intermediates, especially fructose-1,6-bisphosphate (F-1,6-bP)(12), the levels of which are likely significantly lower due to a reduction in glucose-PTS activity. Of note, the enzyme responsible for producing pyruvate (and ATP) by conversion of PEP, pyruvate kinase (*pykF*), is also known to be allosterically activated by F-1,6-bP (35). This drop in Ldh activity could have the effect of increasing the ratio of NADH to NAD^+^ in the *manL* mutant. Perhaps as a means of boosting the levels of NAD^+^, which is needed for glycolysis, the NADH oxidase gene (*nox,* SSA_1127) was upregulated by about 10-fold in the *manL* background. Consistent with the increased flux of pyruvate toward acetyl-CoA, expression of the two genes required for converting acetyl-CoA to acetate and for generation of ATP, phosphate acetyltransferase *pta* along with the rest of the operon SSA_1207 to SSA_1210, which also encode a GTP pyrophosphokinase (RelQ)(36), and acetate kinase (*ackA,* SSA_0192), was increased by 5- and 3-fold, respectively. Biochemical assays showed significantly less lactic acid and more acetate being produced by the *manL* mutant than by SK36 (below). Compared to lactic acid, acetic acid has a higher p*K_a_* value, which may result in less acidification of the cytoplasm and the local biofilm environment.

**Fig. 2.**
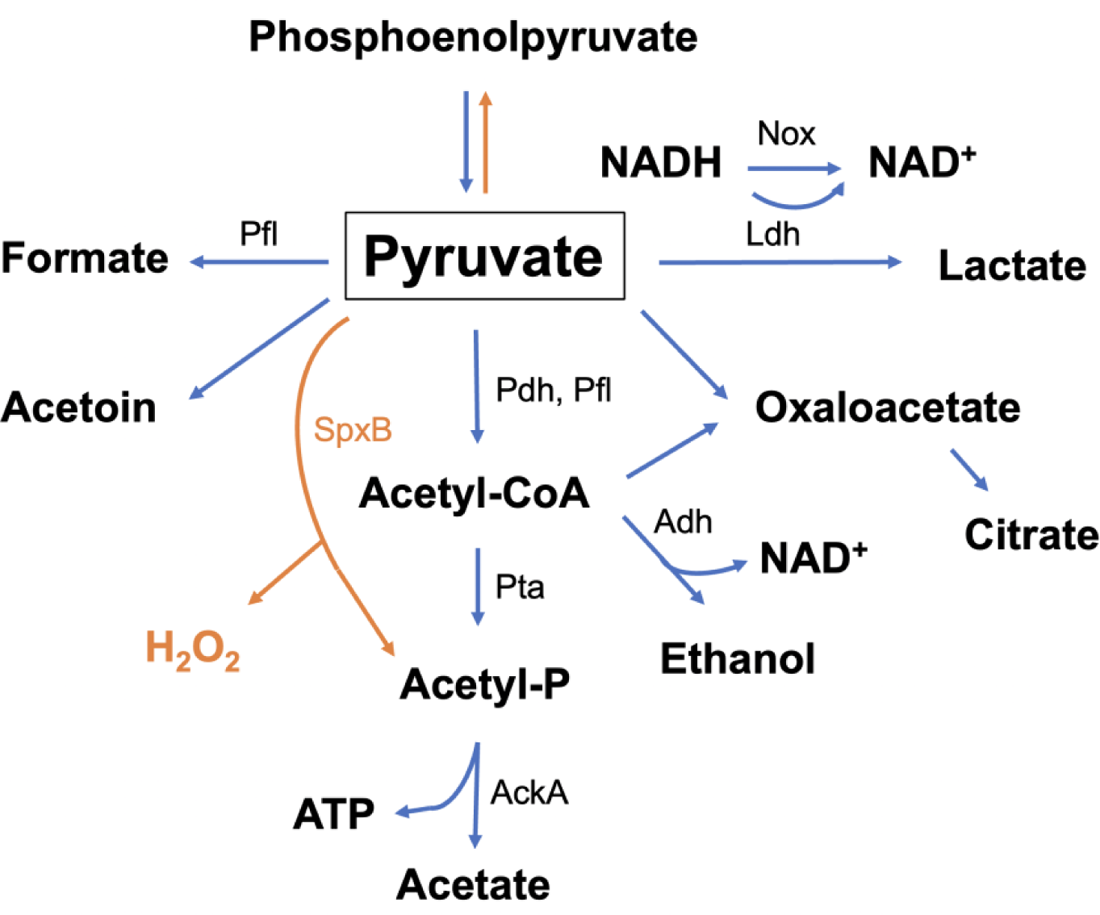
A diagram depicting pyruvate metabolism in streptococci. Phosphoenolpyruvate generated during glycolysis is converted to pyruvate, a central metabolite that can be reduced to lactate with concomitant conversion of NADH to NAD^+^, or further oxidized to acetyl-CoA, formate, acetate, ethanol, acetoin, citrate, etc. Also present in *S. sanguinis* and *S. gordonii* (in orange), but not in *S. mutans*, are a pyruvate oxidase (SpxB) that converts pyruvate into Acetyl-P and H_2_O_2_, and two enzymes capable of performing gluconeogenesis (82). Deletion of glucose-PTS (*manL*) likely results in reduced Ldh activity and enhanced flux toward oxidative branches of pyruvate metabolism that produce H_2_O_2_ and additional ATP molecules. Pfl, pyruvate-formate lyase; Ldh, lactate dehydrogenase; Nox, NADH oxidase; Pdh, pyruvate dehydrogenase; SpxB, pyruvate oxidase; Pta, phosphotransacetylase; AckA, acetate kinase; Adh, alcohol dehydrogenase.

Consistent with the notion that carbon flux away from lactate production toward alternative end products like formate and ethanol, increased mRNA levels were observed for genes encoding enzymes that participate in formate fixation (SSA_0432 to SSA_0435, increased by ∼10-fold), as well as two apparent alcohol metabolizing genes, *adhE* (SSA_0068, increased by 9-fold) and *adhP* (SSA_1917, increased by 3-fold)(37). Alcohol-generating activities of alcohol dehydrogenases may also help to restore the NADH:NAD^+^ balance by oxidizing NADH. The impact of *manL* deletion on acetoin production and metabolism appeared less clear, as the operon predicted as acetoin dehydrogenase, SSA_1174 to SSA_1178 (38) showed a 3-fold increase in expression, as did a putative butanediol dehydrogenase (SSA_0572, increased by 5-fold), yet the *ilv* operon (SSA_1967 to SSA_1971) encoding gene products that convert pyruvate to acetolactate, a precursor to acetoin (or branched-chain amino acids), was downregulated about 2-fold. One plausible explanation for these changes in the *manL* mutant is to deal with accumulation of acetoin, which could be toxic at elevated levels, that resulted from the accumulation of pyruvate (39). Another system frequently affected by PTS activities and changes in intracellular energy status is the *glg* operon (SSA_0775 to SSA_0779), which was upregulated 4- to 7-fold in the *manL* mutant.

#### ii) Carbohydrate transport and degradation

Similar to many lactic acid bacteria, *S. sanguinis* can ferment a variety of carbohydrates (40). Deletion of *manL* in SK36 resulted in enhanced expression of genes for carbohydrate transporters and associated metabolic enzymes. A 24-gene locus, SSA_0508 to SSA_0531, that is predicted to encode enzymes for catabolism of propanediol and ethanolamine (41) was upregulated 2- to 5-fold. This locus is part of a 70-kbp, genomic island-like sequence that also includes enzymes for vitamin B_12_ biosynthesis (33). When ethanolamine was supplied as the sole carbohydrate, both SK36 and its *manL* mutant failed to produce significant growth (data not shown). We cannot rule out the possibility that *S. sanguinis* is capable of catabolizing ethanolamine for energy extraction or for the benefit of ammonia release (below), just not for growth, or that additional factors are needed for growth that may be available under certain conditions.

Also showing increased expression in the *manL* mutant was a 14-gene locus predicted to encode an ascorbate utilization pathway, including genes for a PTS permease for ascorbate (42), degradative enzymes, and a (PTS regulatory domain-) PRD-containing transcription anti-terminator. SK36 grew well with ascorbate as the sole carbohydrate, whereas the *manL* mutant had a modestly slower growth rate (Fig. S2). ManLMNO can transport carbohydrates other than glucose (e.g., mannose, galactose, GlcN, GlcNAc), so the data could be interpreted to mean that ManLMNO can also internalize ascorbate. Third, increased in expression by about 8-fold were genes SSA_0053 to SSA_0062 encoding a β-galactosidase (*bgaC*), a mannose/fructose/N-acetylgalactosamine PTS transporter, a sugar isomerase (*agaS*), a tagatose-bisphosphate aldolase (*lacD*), and a galactose mutarotase (*galM*). A homologous system (*aga* operon) in *S. pneumoniae* was shown to be inducible by N-acetylgalactosamine and likely required for uptake and metabolism of N-acetylgalactosamine-related polysaccharides (43, 44). Another notable DE gene was that encoding a maltose/glucose PTS transporter SSA_0379 sharing homology with the maltose transporter MalT in *S. mutans* (45). SSA_0379 was upregulated by about 47-fold in the *manL* mutant. It is possible that this gene product can compensate for the loss of ManL by transporting glucose.

Lactose/galactose metabolic genes were also affected by deletion of *manL*. The lactose-PTS genes *lacTFEG* were upregulated by about 3-fold. In contrast, the tagatose pathway genes *lacABCD* were down-regulated by 3-fold, while the Leloir pathway (SSA_1008 and SSA_1009) was upregulated by about 3-fold (46). In addition, deletion of *manL* also resulted in elevated expression by multiple predicted extracellular β-galactosidases (SSA_0053 and SSA_0271), an N-acetylneuraminate lyase (SSA_0078), an endoglucanase (SSA_0182), two putative pullulanases (SSA_0453 and SSA_2268), a β-hexosamidase or β-N-acetylhexosaminidase (SSA_1065, increased by 15-fold), and a homologue of fructan hydrolase (*fruA*, SSA_2023)(28, 47). Additional carbohydrate transporters included a maltose/maltodextrin ABC-transporter (SSA_1298 to SSA_1300), a PTS operon (SSA_0219 to SSA_0224) homologous to the fructose/mannose-PTS permease (*levDEFG*) originally identified in *S. mutans* (48), and other predicted carbohydrate transporters: SSA_0074 to SSA_0076, SSA_0268 to SSA_0270, SSA_0456, and SSA_2084, all showing various levels of enhancement in mRNA levels (Table S1). It was clear that the glucose-PTS ManLMNO exerts wide-ranging, negative regulation on a variety of secondary carbohydrate-transport and catabolic genes encoded in the genome of SK36. Reducing ManLMNO-dependent carbohydrate transport may therefore allow the cells to metabolize at a lower rate (than the wild-type growing on glucose), but with a better yield and access to a greatly expanded repertoire of substrates. Modulation of gene expression in this manner may be particularly beneficial for persistence in the oral cavity and during systemic infections. For example, strain SK36/*manL* growing in 0.05% glucose, which is close to normal blood sugar levels, produced a particularly higher yield than the wild type (Fig. S2; OD_600_ at 0.34 vs 0.23).

#### iii) Alkali production

As a relatively acid-sensitive early colonizer of the oral cavity, the ability to generate alkaline compounds enhances the competitiveness of *S. sanguinis*. We have reported that strain SK36/*manL* can persist much better in post-exponential phase cultures and can reach significantly higher final pH values than the wild type in certain media or with added arginine (32). The RNA-Seq analysis showed that deletion of *manL* resulted in elevated expression of several pathways with activities that either release ammonia or reduce the production of acids in the cytoplasm. The arginine deiminase (*arc*) operon, SSA_0736 to SSA_0743, was expressed at levels at least 10-times higher than in wild type (Fig. 1). AD benefits streptococci both via release of ammonia and generation of ATP (49). Genes of the *arc* operon are known to be controlled by CcpA, pH and oxygen tension (50–52). Clearly, though, ManL has a major influence, directly or indirectly, on AD gene expression.

Second, SSA_0429 to SAA_0435 region encodes for a set of enzymes for histidine and formate metabolism. Histidine-ammonia lyase (SSA_0429, up by 4-fold) and formimidoyltetrahydrofolate cyclodeaminase (SSA_0433, up by 15-fold) may release substantial amounts of ammonia, especially given the likely greater availability of formate in the mutant (see below). Similarly, significantly increased expression of both ethanolamine ammonia lyase (above), encoded by SSA_0518 to SSA_0520, and alanine dehydrogenase (SSA_1615, by 13-fold), is expected to increase ammonia production given their respective substrates. Lastly, increased flux through the acetoin dehydrogenase pathway (SSA_1173 to SSA_1178, up by 3-fold) may reduce acidification of the cytoplasm by avoiding accumulation of acidic byproducts.

#### iv) Membrane biosynthesis

Another significant phenotype of the *manL* mutant was its reduced tendency to undergo autolysis (32). A group of peptidoglycan (PG)-degrading enzymes are known to play pivotal roles in autolysis and programmed cell death by bacteria such as *Staphylococcus aureus, S. pneumoniae,* and *S. mutans* (53–55). No genes homologous to these proven PG hydrolase genes were identified among those differentially expressed in the *manL* mutant. However, there were strong indications for enhanced glycerolipid metabolism by the mutant. One apparent operon structure, SSA_1826 to SSA_1828, predicted to encode glycerol uptake and metabolic genes, showed the greatest increase (71- to 91-fold, Fig. 1) in expression among all genes. Also, SSA_0049 to SSA_0051, which encode three dihydroxyacetone kinases, showed 10- to 12-fold increase in mRNA levels. These gene products function in catabolic as well as anabolic metabolism of glycerolipids (56). With the overall carbon flux being directed towards acetyl-CoA, a primary substrate for fatty acid biosynthesis, it is likely that the *manL* mutant could have alterations in membrane biogenesis that enable it to better persist under environmental stressors like low pH and increased H_2_O_2_ (57). Membrane remodeling has been strongly associated with acid tolerance and adaptation in *S. mutans* and a variety of other bacteria (58, 59).

#### v) Attachment and other functions

A recent study identified *S. sanguinis* as the only streptococcal species to possess a type-IV pili (Tfp) gene cluster. Although SK36 is incapable of Tfp-mediated twitching motility, this locus (SSA_2302 to SSA_2318) encodes for biosynthesis of pili that accounted for >40% of attachment to two human epithelial cell lines (60), and were required for invasion of human aortic endothelial cells and pathogenesis in an infective endocarditis model (61). The entire 18-gene *tfp* locus showed about 2-fold increase in mRNA levels due to loss of *manL*, indicative of the ability of glucose-PTS to regulate adherence and colonization in oral cavity. Further, a putative C69-family dipeptidase (SSA_1591) was 21-fold more highly expressed in the *manL* background. SSA_1591 is a homologue of the *S. gordonii* amylase-binding protein (AbpB) (62, 63), which binds to human salivary amylase and also enhances the activity of the glucosyltransferase GtfB of *S. mutans* (64); both of these activities have proven roles in bacterial attachment and biofilm development.

Opposite to its effect on the *arc* operon, deletion of *manL* significantly downregulated the expression of several arginine biosynthetic genes, including SSA_2141 and SSA_2142 (by 7-fold), and SSA_0757 to SSA_0760 (by 3-fold). Combined with greater AD activities, this could result in a significant reduction or depletion of intracellular arginine. A significant portion of the genes showing lower expression in the *manL* mutant, totaling 34, are considered as “hypothetical proteins” or having “unknown function”, while only 15 such uncategorized genes are present in the upregulated group.

#### vi) Comparison with the CcpA regulon

A recent study (23) delineated the CcpA regulon in *S. sanguinis* by culturing SK36 and its otherwise-isogenic *ccpA* mutant on the glucose-containing, rich medium BHI. By applying RNA-Seq analysis, this study identified a total of 169 DE genes, primarily affecting pathways for central metabolism, utilization of carbohydrates and amino acids, and the PTS. Loss of *ccpA* also resulted in additional phenotypes including overproduction of H_2_O_2_ and excretion of pyruvate (21, 23, 65). While a deficiency in glucose-PTS is expected to cause reduced carbon flux and relief of CCR in glucose-containing medium, we observed limited overlap (22%) of genes affected by deletion of *ccpA* and by deletion of *manL*. Part of this difference might have resulted from the experimental conditions under which these studies were conducted: the CcpA study used anaerobically grown cells whereas we used an aerobic incubator containing 5% CO_2_. We have denoted in our DE genes those identified by the CcpA study and those by a separate bioinformatic analysis on the CcpA regulon (23)(Table S1). Significantly, we counted a total of 71 genes that belonged to the CcpA regulon yet showed no significant change, or had changes in expression in the opposite direction, in the *manL* mutant. The latter group included SSA_2096, SSA_2249, and SSA_2318 (Table S1). Furthermore, deletion of *manL* resulted in a 3-fold increase in expression of *ccpA* itself, whereas the CcpA study also identified *manL* as being negatively regulated by CcpA (23). Together these findings demonstrate that the ManLMNO permease in *S. sanguinis* plays a dominant role in governing central metabolism and metabolism of many secondary carbohydrates, and that part of the effects on gene expression occur indirectly through CcpA. The rest of this study was devoted to understanding the metabolic and fitness impacts of glucose-PTS-mediated regulation to *S. sanguinis* and related bacteria.

### Targeted metabolomics of organic acids indicated altered metabolism by the *manL* mutant

*S. sanguinis* under aerobic laboratory conditions undergoes a mixed-acid fermentation, producing a variety of organic acids. To assess the role of the ManLMNO permease in carbohydrate-mediated regulation, we harvested bacterial cells from TY-glucose cultures and measured the abundance of 8 common organic acids using liquid chromatography coupled with mass spectrometry (LC-MS/MS). The results (Fig. 3), though not statistically significant, showed substantially reduced lactic acid and increased pyruvate levels in bacterial cells of the *manL* mutant, which is consistent with measurements previously made in culture supernates using biochemical assays (32). The *manL* mutant also produced significantly less fumarate, despite the fact that the SK36 genome apparently lacks the genes to produce fumarate (66). Interestingly, the metabolomic analysis showed no detectable levels of citrate and only small amounts of α-ketoglutarate.

**Fig. 3.**
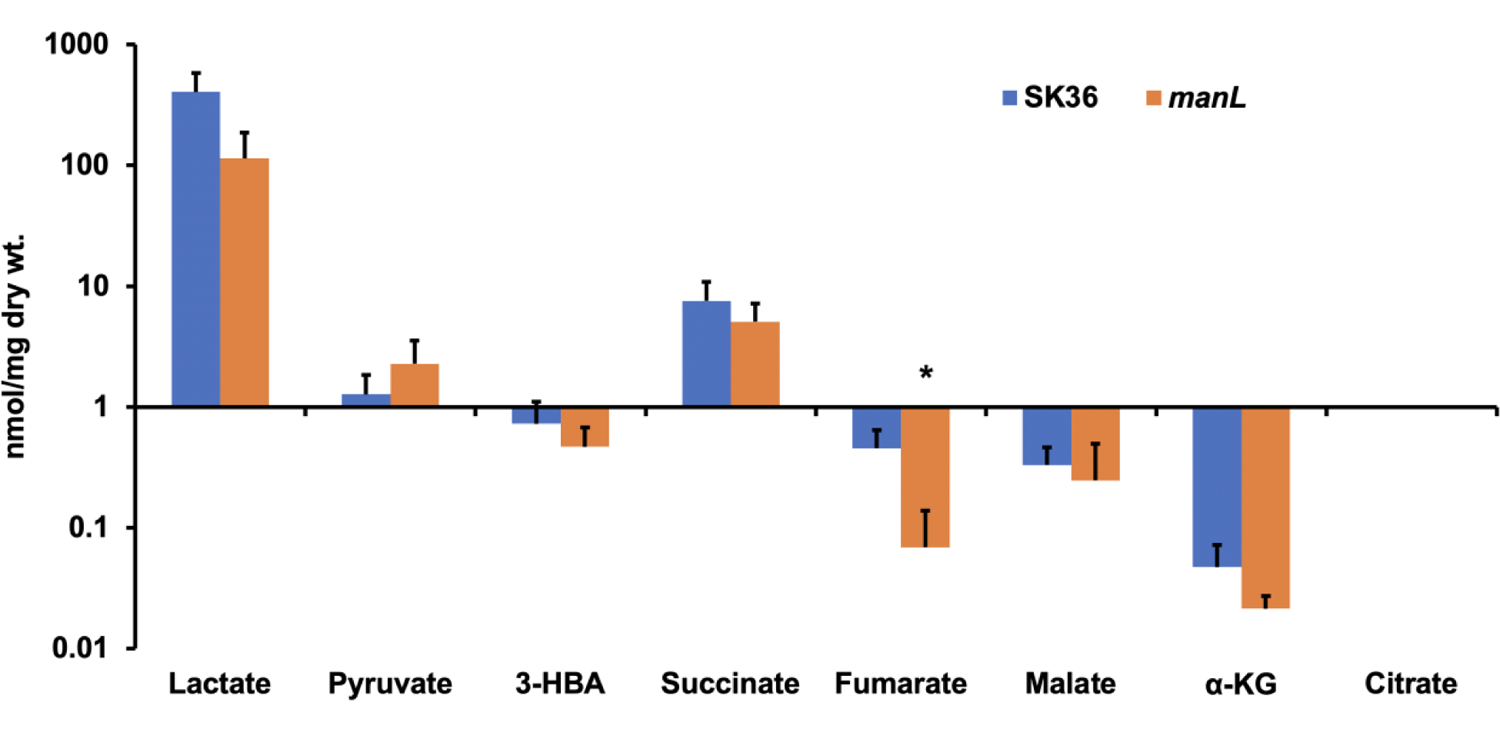
Targeted metabolomics of 8 organic acids. SK36 and SK36/*manL* were cultivated overnight in BHI medium and sub-cultured in TY-glucose after a 20-fold dilution. Bacterial cells from exponential-phase cultures were harvested by centrifugation, from which metabolites were extracted for liquid chromatography and mass spectrometry. The data were derived from 3 independent cultures (n = 3) and the results were assessed for statistical significance using Student’s *t*-test (*P* value: * <0.05). 3-HBA, 3-hydroxybutyric acid; α-KG, α-ketoglutarate.

To complement the metabolomic analysis, we also employed individual biochemical assays to measure the levels of lactate, acetate, formate, citrate, and acetoin in the supernates of strains of SK36, the *manL* mutant, and the *ccpA* mutant (Table 1). TV-glucose medium was used instead of TY-glucose to avoid contaminating metabolites originated from yeast extract. As shown in Table 1, deletion of *manL* in SK36 resulted in a 4.0-fold reduction in lactate levels, but significantly higher acetate (1.8-fold), citrate, and especially higher levels of formate (6.9-fold) in the supernates. There was also a 10% reduction in total acids in the *manL* mutant compared to the wild type. In contrast, deletion of *ccpA* resulted in significantly higher production of all four acids and a 78% increase in total acids. No acetoin was detected (detection limit ∼50 μM; data not shown) in cultures prepared with TV-glucose, TY-glucose, or BHI. Also, different from results obtained with cells grown in TY-glucose or FMC medium, but consistent with findings from *S. mutans* (67), no significant amounts of pyruvate were detected in cells grown in TV-glucose. Thus, SK36/*manL* yielded a profile of organic acids that was consistent with the reprogramming of pyruvate metabolism revealed by RNA-Seq, i.e. increased flux toward pyruvate-formate lyase, pyruvate dehydrogenase, and pyruvate oxidase with a concomitant decrease in lactate dehydrogenase activity (Fig. 2). A reduction in total acids also supported the notion that the *manL* mutant undergoes a more efficient mode of pyruvate metabolism, as production of acetate yields an additional ATP compared to production of lactate. Increased acid production by the *ccpA* mutant could be a result of elevated expression of the glucose-PTS and a boost to the carbon flux through glycolysis and pyruvate metabolism (20), although no difference in mRNA levels of *ldh* or *pykF* was reported in the *ccpA* mutant (23). Since lactic acid is stronger than acetic acid, but weaker than formic acid, the net effect of these changes appeared consistent with, albeit insufficient to explain completely, the higher final pH values of SK36/*manL* cultures. Rather, enhanced alkali production accounts for the final pH achieved by the *manL* mutant. In addition to AD, activation of the aforementioned alkali-generating activities requires further study to assess their relative contributions to pH homeostasis and fitness. Furthermore, the resting pH measured in SK36 glucose cultures (approximately 4.70)(32) was above the p*K_a_* values of lactic and formic acids, however slightly below that of acetic acid (4.76); the pH of SK36/*manL* (5.25)(32) was above the p*K_a_* values of all three acids. Therefore, a greater proportion of the acetic acid should be protonated in SK36 culture supernates than in those of SK36/*manL*. Since the bacterial membrane is much more permeable to protonated acids than their anionic form, the wild type is thus posited to have higher cytoplasmic levels of acetate than the *manL* mutant, which is less favorable in terms of membrane proton force (ΔpH) and could trigger autolysis, similar to acetate-dependent programmed cell death in certain other Gram-positive bacteria (68).

**Table 1.** Organic acids measured in the supernates of 20-h TV-glucose cultures. Each strain was represented by 3 biological replicates and the measurements were normalized against their optical density and presented as average and standard deviation (mM/OD_600_). Asterisks denote statistical significance assessed by Student’s *t*-test (*P* values: * <0.05, ** <0.01, *** <0.001) when compared to results obtained with the wild type.

| Strains | Lactate | Acetate | Formate | Citrate | Total acid |
| --- | --- | --- | --- | --- | --- |
| *S. sanguinis* |  |  |  |  |  |
| SK36 | 14.36 ± 0.65 | 2.68 ± 0.25 | 1.14 ± 0.12 | 0.04 ± 0.007 | 18.22 |
| *manL* | 3.62 ± 1.27*** | 4.92 ± 0.51** | 7.90 ± 0.93*** | 0.07 ± 0.003* | 16.51 |
| *ccpA* | 26.47 ± 1.82** | 4.03 ± 0.72 | 1.92 ± 0.49*** | 0.09 ± 0.014 | 32.51 |
| *S. mutans* |  |  |  |  |  |
| UA159 | 30.50 ± 4.30 | 1.85 ± 0.15 | 5.17 ± 0.26 | 0.04 ± 0.04 | 37.55 |
| *manL* | 13.31 ± 1.75** | 4.37 ± 0.32*** | 12.32 ± 0.92*** | 0.05 ± 0.02 | 30.05 |
| *S. gordonii* |  |  |  |  |  |
| DL1 | 32.07 ± 1.79 | 0.30 ± 0.11 | 0.46 ± 0.11 | 0.02 ± 0.003 | 32.85 |
| *manL* | 28.00 ± 1.69* | 1.05 ± 0.29** | 1.17 ± 0.06*** | 0.04 ± 0.011 | 30.26 |

### Two-species biofilms highlight the role of glucose-PTS in bacterial fitness and competition

A two-species biofilm model was utilized to study the impact of deletion of *manL* on the competitiveness of *S. sanguinis* when co-cultured with *S. mutans* UA159 under various environmental conditions. In these experiments, SK36 or SK36/*manL* was allowed to form biofilms on a saliva-coated glass surface for 1 day, followed by inoculation with UA159. The biofilms were incubated for 2 additional days then CFU were enumerated. When the standard BMGS medium (69) was used, UA159: SK36 biofilms yielded nearly 2-log higher CFU of *S. sanguinis* than that of *S. mutans*. Deletion of *manL* in SK36 resulted in enhanced recovery of *S. mutans*, such that more than 1-log higher CFU of UA159 were present in the UA159: SK36/*manL* biofilms than the UA159: SK36 biofilms (Fig. 4A). During some of the repetitions, the UA159: SK36/*manL* group also returned significantly greater *S. sanguinis* counts than did the group UA159: SK36 (Fig. 4B and Fig. S3), indicative of enhanced fitness of the *manL* mutant.

**Fig. 4.**
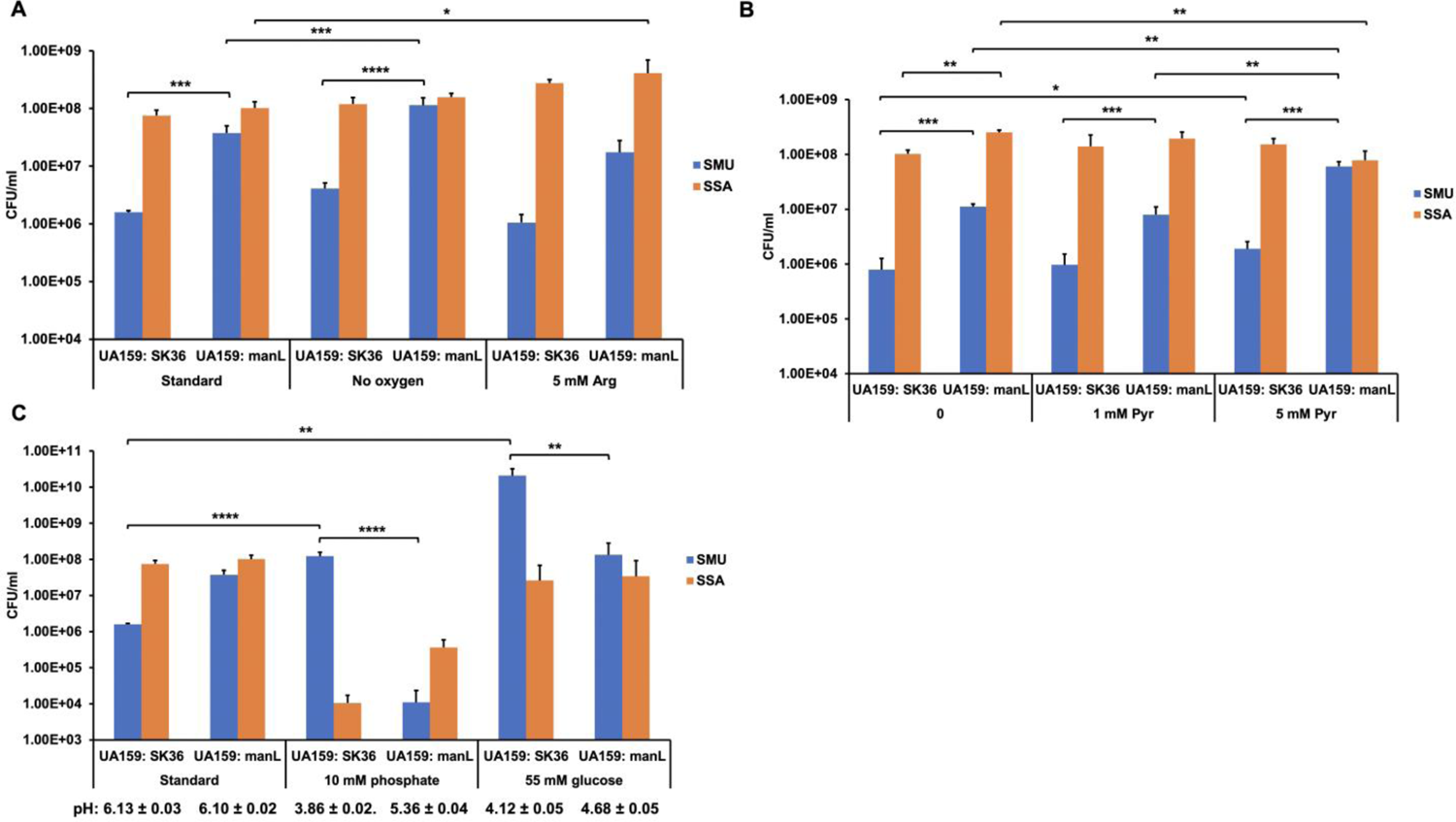
Two-species biofilm analysis. *S. sanguinis* SK36 and *manL* mutant (n = 4) carrying a chromosomally borne, selectable antibiotic resistance marker were used to inoculate a biofilm on saliva-coated glass surfaces. *S. mutans* was introduced 1 day later. After two additional days of cultivation, biofilms were harvested for CFU enumeration. The growth medium and atmosphere of the biofilm cultures were, when specified, modified to test the effects of (A) oxygen and arginine (Arg), (B) extracellular pyruvate (Pyr), and (C) buffer capacity and carbohydrate availability. The results of each experiment are presented as averages and standard deviations (error bars). Each comparison was assessed for statistical significance by applying two-way ANOVA (*P* values: * <0.05, ** <0.01, *** <0.001, and **** <0.0001).

Given the importance of oxygen metabolism and pH homeostasis in the interactions between *S. sanguinis* and *S. mutans*, we tested the effects of oxygen or arginine in the medium. When biofilms were grown as above in an anaerobic jar, slightly more *S. mutans* cells were recovered in the UA159: SK36 group compared to the same group cultured in an ambient incubator containing 5% CO_2_. In contrast, the UA159: SK36/*manL* group returned even more *S. mutans* cells (Fig. 4A) than the UA159: SK36 group, likely attributable to the improved H_2_O_2_ production of the *manL* mutant inhibiting the growth of and/or killing *S. mutans*. Conversely, when 5 mM of additional arginine was present in the medium, a slight reduction in *S. mutans* counts was noted in both groups, though not statistically significant. Concurrently, arginine addition promoted the persistence of *S. sanguinis,* especially the *manL* mutant (Fig. 4A). This result was consistent with increased expression of the arginine deiminase system in the *manL* mutant moderating acidification of the *S. sanguinis* cytoplasm and the environment (4).

We have reported that strain SK36/*manL* excretes significantly more pyruvate into the medium than the wild type, a substrate that can be internalized and metabolized by *S. mutans* through a specialized transporter (70). To investigate if pyruvate released by SK36/*manL* benefited UA159, pyruvate was added into the medium at physiologically relevant concentrations (Fig. 4B). When present at 1 mM, exogenous pyruvate showed no impact on the CFU recovered of either bacterium. At 5 mM, exogenous pyruvate significantly improved recovery of *S. mutans*, especially when the *manL* mutant of SK36 was used. This result supported the notion that *S. mutans* benefits from the additional pyruvate excreted by the *manL* mutant of SK36, compared to that produced by wild-type SK36.

To better mimic a cariogenic environment, we reduced the buffer capacity and significantly increased the glucose content of the medium. The buffer capacity of human saliva ranges from 5 to 24 mM HCO_3_^-^ and 3 to 5 mM phosphate, with an average of these buffers being around a total of 10 mM (71, 72). As shown in Fig. 4C, when the total phosphate levels were reduced from 72 mM in the standard BMGS to 10 mM, UA159: SK36 biofilm showed a dominance by *S. mutans* and a drastically lower abundance of *S. sanguinis*. Interestingly, UA159: SK36/*manL* biofilm showed the opposite pattern, with *S. sanguinis* outcompeting *S. mutans*, although both species returned significantly lower CFU than in the standard BMGS. When the glucose content was increased from 18 mM to 55 mM (Fig. 4C), *S. mutans* expectedly showed a substantial increase in abundance when co-cultured with SK36. However, this benefit was all but eliminated in the presence of SK36/*manL.* When the pH of the culture was measured at the end of each experiment, the results were consistent with the observed trends of bacterial abundance. Specifically, at 10 mM phosphate, the UA159: SK36 biofilm had a final pH of 3.86 while the UA159: SK36/*manL* group had a final pH around 5.36; and they produced a pH around 4.12 and 4.68, respectively, when growing on 55 mM glucose. Since the *manL* mutant was likely more acid tolerant and expressed higher AD activities than the wild type, these differences in final pH were probably due to arginine catabolism that resulted in a higher end-point pH in biofilms containing SK36/*manL*.

*S. mutans* has little or no capacity to elicit a pH rise. Clearly, then, these biofilm studies support that the glucose-PTS plays an important role in the ability of *S. sanguinis* to cope with the rapid acidification of dental plaque due to the consumption of carbohydrates by negatively regulating alkali production, as the reduction of pH generally coincides with the exhaustion of carbohydrates thus relief of PTS-mediated catabolite repression. As an abundant member of the dental plaque, the ability of *S. sanguinis* to regulate central metabolism and alkali production in response to carbohydrate availability could impact the entire microbiome. Finally, under *in vivo*, cariogenic conditions where the microbiome is exposed to frequent influx of large amounts of fermentable carbohydrates, or insufficient buffer capacity due to certain physiological conditions that restrict salivary flow rate, commensals face the selective pressure for greater acid tolerance and for enhanced bioenergetic efficiency. Under such conditions, mutants with enhanced competitiveness, such as those of the glucose-PTS, could become prominent in the microbiome populations thus contributing to the elevated aciduricity of the biofilm. Further investigation is needed to identify such mutations *in vivo* and understand their significance to pH homeostasis, microbial ecology, and biofilm pathogenicity.

### Deletion of *manL* in *S. mutans* and *S. gordonii* alters bacterial metabolism and competition

*manL* deletion mutants of *S. mutans* UA159 and *S. gordonii* DL1 were utilized to study the effects of such mutations on their metabolism and fitness, given their importance in oral microbial ecology. Quantification of 4 organic acids in planktonic cultures of both mutants showed reduced lactate and increased acetate and formate, relative to the wild type (Table 1). These changes in acid profile echoed the effects of the *manL* deletion in *S. sanguinis* background, although the magnitude of change in *S. gordonii* appeared significantly smaller than in the other two species.

Twenty-hour, TY-glucose planktonic cultures of three wild-type strains and their respective *manL* mutants were assayed for their resting pH, final optical density (OD_600_), CFU/ml, release of eDNA (RFU measurement after binding a fluorescent dye), and excretion of pyruvate. This time point was selected to represent approximately the start of the stationary phase for all 6 strains and to allow sufficient expression by genes normally suppressed by catabolite repression. As shown in Fig. 5, deletion of *manL* in *S. gordonii* strain DL1 resulted in increased final yield in CFU counts, though not in optical density, and reduced eDNA levels. It appeared that DL1/*manL* had reduced autolytic activity, as seen in SK36/*manL* (32). The cultures of DL1/*manL* also showed significantly higher final pH (Fig. 5D) than the wild type and enhanced excretion of pyruvate (Fig. 6A). By contrast, a *manL* mutant of *S. mutans* UA159 did not show any notable difference in accumulation of eDNA or optical density from the parental strain UA159, instead showed a >1-log reduction in CFU (Fig. 5). At the same time, UA159/*manL* (Table 1) produced 2.3-fold less lactic acid compared to the wild type and had a 20% reduction in total acids. However, the mutant also had a 2.4-fold increase in both acetic and formic acid. These results were consistent with the fact that UA159/*manL* did not show a significant increase in the resting pH from the wild type in stationary cultures (around 4.43, compared to 4.41 for the wild type; Fig. 5D); in part because *S. mutans* lacks significant alkali-generating capacity. Likewise, a previous study reported that a *manL* mutant of UA159 was less capable of lowering the pH and tolerating acid byproducts than the wild type in an *in vitro* pH drop assay (73).

**Fig. 5.**
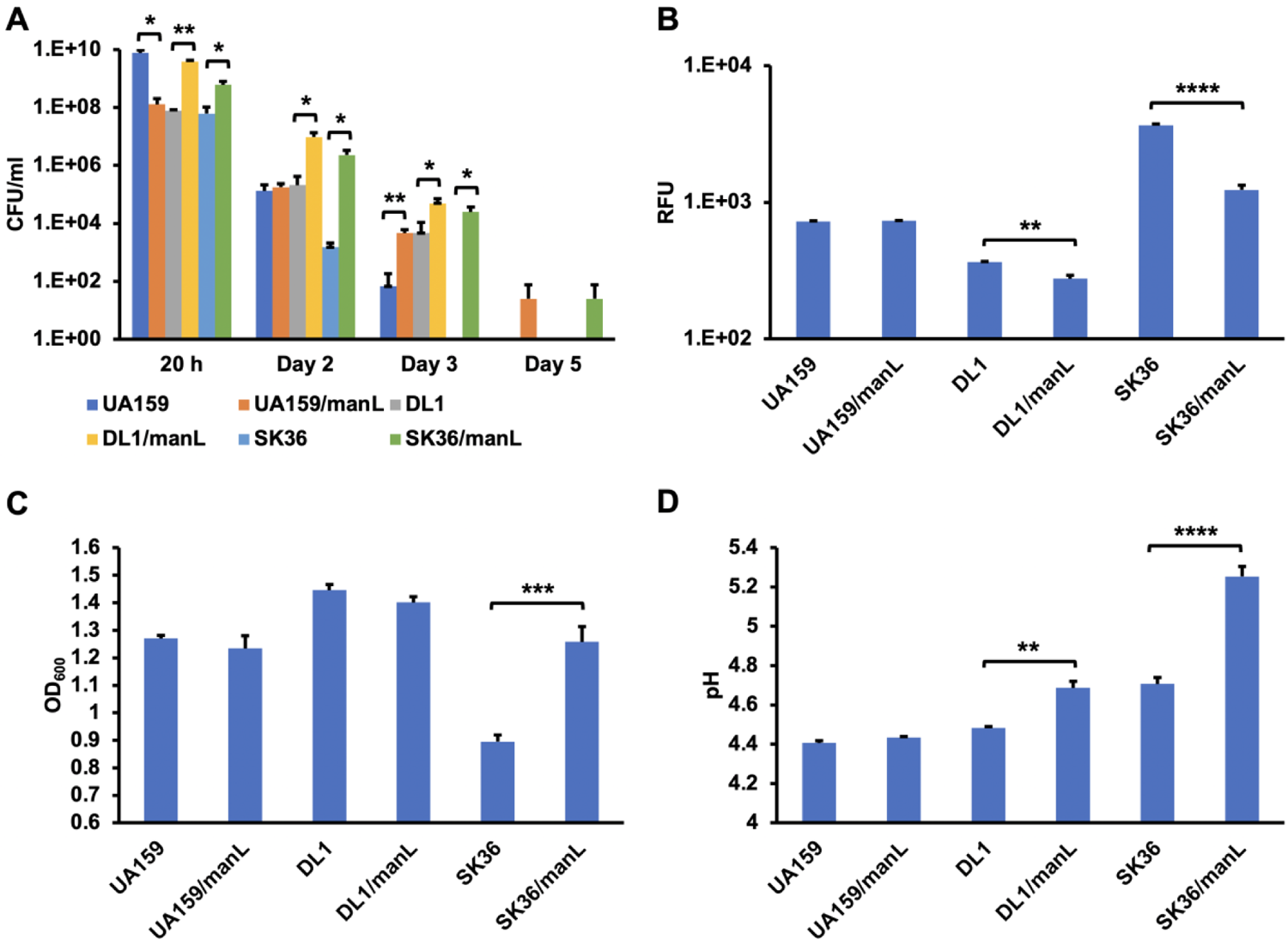
Phenotypic assessment of 3 streptococci and their *manL* mutants. *S. sanguinis, S. gordonii,* and *S. mutans* strains were each (n = 3) cultivated for 20 h (B, C, D) or 5 days (A) in TY medium containing 20 mM of glucose. Each culture was then analyzed at specified time points for A) CFU enumeration, B) eDNA as relative fluorescence units (RFU) resultant of a reaction with a DNA-specific fluorescent dye, C) final OD_600_, and D) resting pH. The results are presented as average and standard deviation (as error bars). Asterisks represent statistical significance between the wild type and its mutant, assessed by Student’s *t*-test (*P* values: * <0.05, ** <0.01, *** <0.001, and **** <0.0001).

**Fig. 6.**
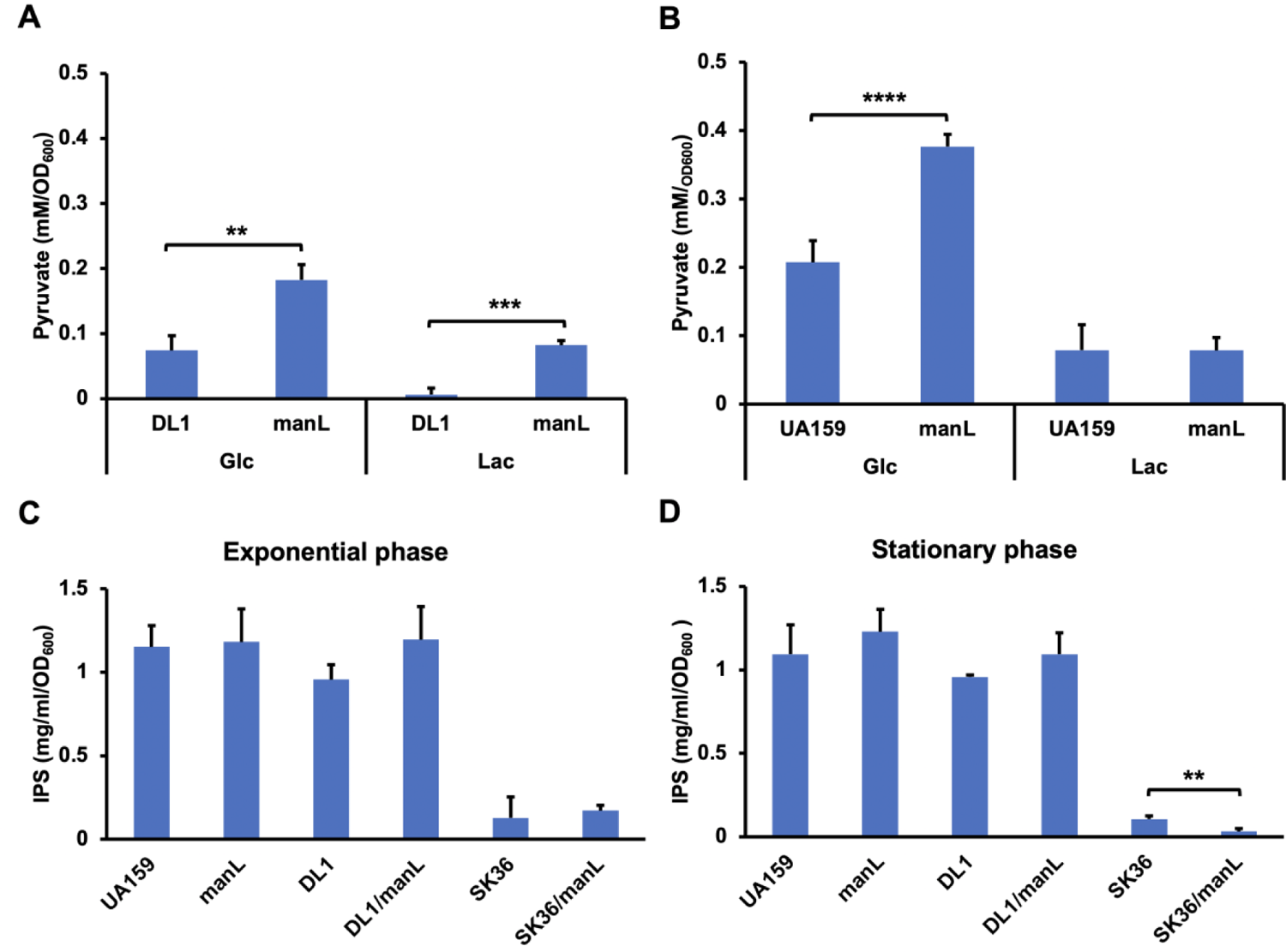
Extracellular pyruvate and IPS measurements of 3 streptococci and their *manL* mutants. (A, B) Extracellular pyruvate in 20-h cultures prepared with TY containing 20 mM glucose or 10 mM lactose. See our previous work for results in *S. sanguinis* (32). (C, D) IPS from exponential and stationary phases of TY-glucose cultures. The data were derived from 3 biological replicates and the results are presented as averages and standard deviations. Asterisks represent statistical significance assessed by Student’s *t*-test (*P* values: * <0.05, ** <0.01, *** <0.001, and **** <0.0001).

Nonetheless, UA159/*manL* did produce significantly more extracellular pyruvate (Fig. 6B) than the wild type. Thus, deletion of glucose-PTS likely allowed the accumulation of more extracellular pyruvate in all three species, although the mutant of DL1 excreted more pyruvate in both glucose- and lactose-based cultures than its parent (Fig. 6A), while UA159/*manL* and SK36/*manL* did so only on TY-glucose [Fig. 6B and (32)]. We also monitored the long-term persistence of three *manL* mutants by incubating their cultures under starvation conditions for 6 days. The (Fig. 5A) *manL* mutants of two commensal species persisted significantly better than their respective parents (Fig. 5A).

IPS synthesis and catabolism has critical functions in streptococcal persistence under starvation conditions (74) and is heavily influenced by CcpA and available carbohydrate source(s) [Table S1 and (20, 30, 75)]. To assess the contribution of IPS to bacterial persistence in glucose-PTS mutants, we measured IPS accumulation in bacterial cells, both during exponential growth and in stationary phase. The overall results (Fig. 6CD) showed no significant difference in IPS levels associated with deletion of *manL* in any of the three species, although SK36/*manL* appeared to have less IPS than the wild type in stationary phase. Therefore, IPS metabolism was not markedly affected by the deletion of glucose-PTS. Although likely induced at the transcription level by relief of CCR, IPS synthesis can be controlled at the enzymatic level in response to high-energy metabolic intermediates such as glucose-6-phosphate, F-6-P, and F-1,6-bP (76), which are likely to be lower in the *manL* mutant due to reduced sugar transport.

We previously reported that SK36/*manL* presented a sugar-specific phenotype in H_2_O_2_ production and antagonism of *S. mutans* (32), producing prominent effects when growing in sugars likely transported by ManLMNO (e.g., glucose, galactose, GlcN, and GlcNAc), but not on lactose. Similar assays were performed here using mutant DL1/*manL* on TY-agar plates supplemented with various carbohydrates, and H_2_O_2_ production was assessed by use of Prussian blue agar plates. A DL1/*ccpA* mutant strain was included for comparison due to its inability to suppress UA159 despite enhanced release of H_2_O_2_ (65). The results (Fig. 7A) showed higher amounts of H_2_O_2_ being secreted by the *manL* mutant than the wild type on glucose, galactose, GlcN, and GlcNAc plates. The DL1/*ccpA* strain showed higher H_2_O_2_ production than wild type and DL1/*manL* on glucose or lactose. Compared to DL1, DL1/*manL* showed enhanced antagonism of *S. mutans* UA159 when grown on glucose, galactose, GlcN, GlcNAc, or both glucose and galactose (Fig. 7B). Therefore, deletion of *manL* and *ccpA* in the *S. gordonii* background elicited comparable peroxigenic phenotypes to that observed in corresponding mutants in *S. sanguinis* SK36 background, although SK36/*manL* produced more H_2_O_2_ than SK36/*ccpA* on glucose (32). Similar to what was reported on glucose and what we observed in *S. sanguinis* background, deletion of *ccpA* in DL1 had no impact to its ability to inhibit UA159 on any of these sugars (32, 65). Together, these results demonstrated the importance of PTS-specific regulation in H_2_O_2_-dependent antagonism of *S. mutans* by both *S. sanguinis* and *S. gordonii*.

**Fig. 7.**
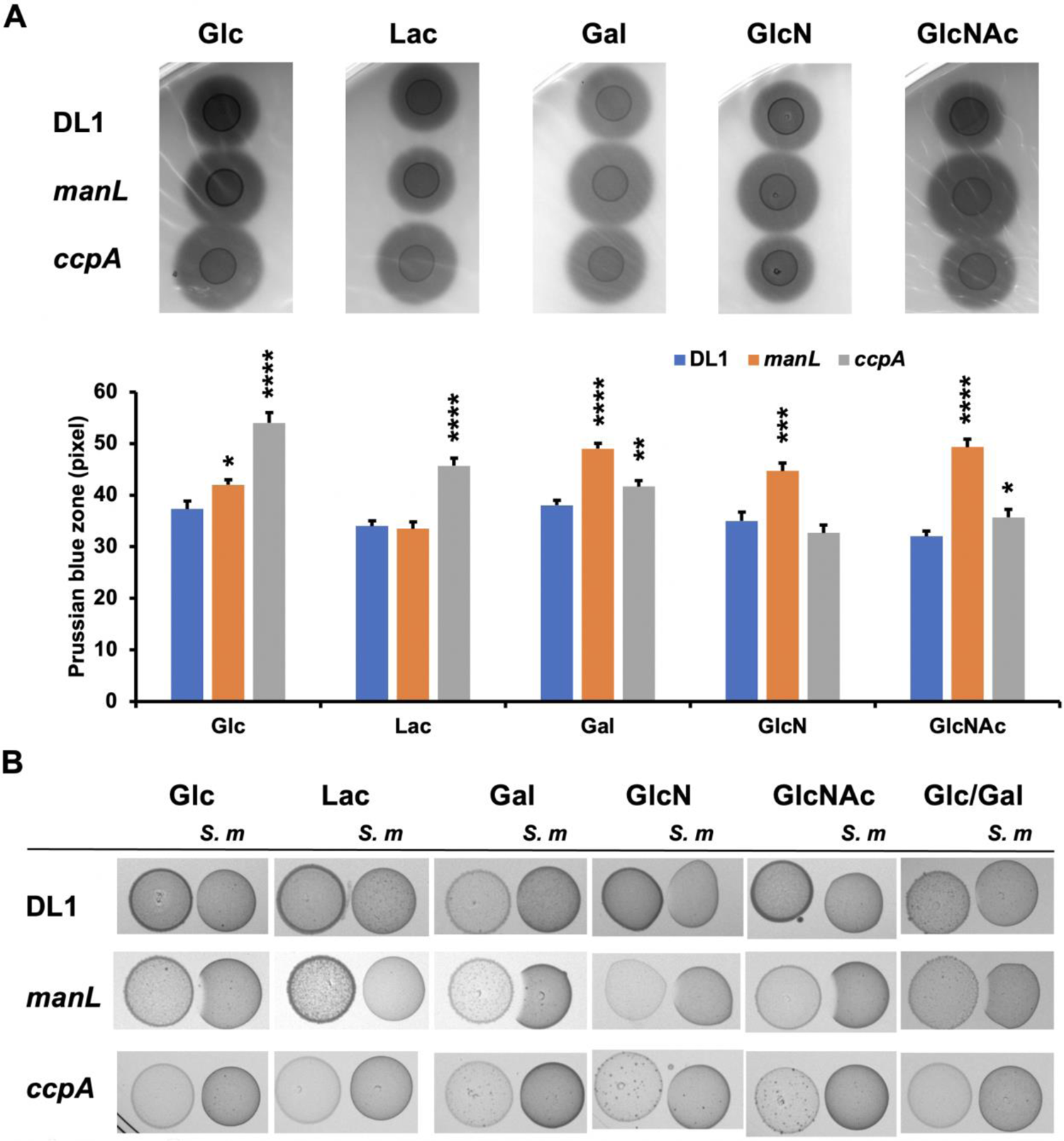
H_2_O2 production (A) and antagonism of *S. mutans* (B) on plates. *S. gordonii* DL1 and its mutants were cultured in TY containing 20 mM of specified carbohydrates till exponential phase, 6 µl of which was dropped onto the agar plates containing respective carbohydrates and incubated for 24 h in an ambient incubator maintained with 5% CO_2_. (A) For measurement of H_2_O_2_ production, the plates included substrates for Prussian blue development (see Methods section for details). Relative production of H_2_O_2_ is represented by the measurements of the width of the blue zone. Statistical significance was assessed using one-way ANOVA (*P* values: * <0.05, ** <0.01, *** <0.001, and **** <0.0001). (B) For antagonism of *S. mutans*, *S. mutans* culture was placed to the right of the colonies. The plates were incubated for another day before photographing. A representative set of graphs are presented.

### Concluding remarks

Streptococci depend on carbohydrates for energy generation and growth. Our transcriptomic, metabolomic, growth, and fitness analyses depicted a metabolic shift in *manL* mutants that traded diminished growth rate for increases in yield and fitness, a strategy that often better supports the stability and diversity of the population (77). Conversely, rapid sugar fermentation by cariogenic bacteria represents the antithesis of such a yield-over-rate strategy and comes with overproduction of acids that drives microbial dysbiosis and dissolution of tooth minerals. It may be argued that microbiome dysbiosis of dental biofilms is primarily a form of metabolic dysregulation at the population level. We hypothesize that distinct populations of the same species exist in the oral cavity by evolving into multiple metabolic variants that are separated by their acid profile, alkali generation, and/or peroxigenic activities, each contributing in different capacities to the pH homeostasis of the microbiome.

Acid adaptation in streptococci occurs by various mechanisms (58), including active proton extrusion by F-ATPase (78), fatty acid remodeling in plasma membrane (59), alteration in PTS activities and specificity (79), alteration in acid end-products (80), alkali generation (4), and acid trapping by glucan matrix (81), etc. It is likely that the glucose-PTS encoded by ManLMNO plays different roles in gene regulation and physiology concerning aciduricity of *S. mutans* vs. two commensal streptococci. Although deletion of *manL* triggered a reprogramming of pyruvate metabolism in all three species, including similar changes in their acid end-product profile, the impact of loss of ManL on pH homeostasis was clearly more significant in the two commensals (especially *S. sanguinis*) than in *S. mutans*; likely due to the conservation of various alkali-generating capabilities in the commensals and their regulation by ManLMNO. This divergence in the role of ManLMNO is especially significant concerning competitions between commensals and cariogenic pathobionts, as the latter are uniquely proficient in fermenting sugars and releasing acids under low-pH conditions. Furthermore, a recent *in silico* study highlighted another difference in metabolic potentials between *S. sanguinis* and *S. mutans* that may contribute to their competitiveness (82)(Fig. 2): *S. sanguinis*, as well as *S. gordonii,* maintains two additional genes (SSA_1053 and SSA_1012) that may enable it to conduct gluconeogenesis from pyruvate. If so, the *manL* mutants of these commensals could possess a unique fitness advantage over *S. mutans* by undergoing gluconeogenesis during starvation (Fig. 5A), especially considering the elevated levels of pyruvate in these mutants. Conversely, *S. mutans* may be able to internalize and utilize the pyruvate secreted by commensals due to its unique pyruvate transporter (LrgAB)(70). However, this benefit may be limited by the availability of carbohydrates, as the source of pyruvate, and oxygen levels which favor H_2_O_2_ production. Last, *S. gordonii* is similar to *S. sanguinis* in composition of core genomes and metabolism (83), but our study indicated that the glucose-PTS ManLMNO in *S. gordonii* may have a significantly different influence on metabolism and fitness than it does in *S. sanguinis*, chiefly in organic acid profile (Table 1, Fig. 6A) and pH homeostasis (Fig. 5D). Research is underway to unravel the complex regulation exerted by *manLMNO*-encoded glucose-PTS in these model organisms and to understand its impact on microbial ecology in dental microbiome.

## Materials and Methods

### Bacterial strains and culture conditions

*S. sanguinis* SK36, *S. mutans* UA159, *S. gordonii* DL1 (Table 2), and their respective isogenic mutants were maintained on BHI (Difco Laboratories, Detroit, MI) agar plates containing 50 mM potassium phosphate buffer (pH 7.2) and used within a week. Antibiotics were added to the agar, when necessary, at the following concentrations: kanamycin (Km), 1 mg/ml; erythromycin (Em), 10 μg/ml; and spectinomycin (Sp), 1 mg/ml. A Tryptone Yeast-extract (TY) medium supplemented with specified carbohydrates was routinely used as the medium for assays without addition of any antibiotics. TY agar was used for plate-based competition assays, and modified to measure bacterial secretion of H_2_O_2_ by inclusion of FeCl_3_·6H_2_O and potassium hexacyanoferrate (III) (84). A Tryptone-Vitamin (TV) medium (85) was used for certain biochemical assays to avoid contaminating metabolites from yeast extract. Biofilms were formed on a glass surface using a biofilm medium [BM, (86)] modified to possess different buffer capacities (72 mM or 10 mM total phosphate) or specified amounts of carbohydrates and other compounds. Unless specified otherwise, all cultures were incubated in a 37°C, ambient-atmosphere incubator supplemented with 5% CO_2_.

**Table 2.** Strains used in this study.

| Strains | Relevant characteristics<sup>a</sup> | Source or reference |
| --- | --- | --- |
| MMZ1896 | *S. sanguinis* wild type SK36 | Kitten laboratory |
| MMZ1616 | SK36 *manL::Em* | MMZ1896 (32) |
| MMZ1617 | SK36 *manL::Km* | MMZ1896 (32) |
| MMZ1945 | SK36 *manL<sup>+</sup> gtfP::Em* | MMZ1896 |
| MMZ2003 | SK36 *manL::Sp gtfP::Em* | MMZ1945 |
| MMZ1913 | SK36 *ccpA::Km* | MMZ1896 (32) |
| DL1 | *S. gordonii* wild type | ATCC 49818 |
| MMZ898 | DL1 *manL::Km* | DL1 (28) |
| MMZ877 | DL1 *ccpA::Sp* | DL1 (28) |
| UA159 | *S. mutans* wild type, *perR*<sup>+</sup> | ATCC 700610 |
| MMZ2020 | UA159 *manL::Km* | From UA159 |

*Em*, *Km*, and *Sp*: resistance against erythromycin, kanamycin, and spectinomycin, respectively.

### Construction and phenotypic characterization of glucose-PTS mutants

Genetic mutants were engineered using an allelic exchange strategy performed by natural transformation of streptococcal cultures with ligation products of DNA fragments containing homologous flanking sequences, and in place of the gene of interest, genetic cassettes encoding resistance against antibiotics Km, Em, or Sp (32, 87). Primers used in this process are included in Table S2. Naturally competent bacterial cultures were obtained using competence-stimulating peptide unique to each species at levels specified in previous publications (*S. mutans* and *S. sanguinis*)(88, 89), or by adding horse serum (*S. gordonii*)(90). DNA fragments necessary for these experiments were generated by PCR and ligated using a Gibson Assembly reaction as detailed elsewhere (32). All genetic mutants were validated by Sanger sequencing targeting the manipulated region of the genome.

Planktonic cultures of various bacterial strains were prepared by diluting (5000-fold) their overnight BHI cultures with TV or TY media supplemented with specified carbohydrates and incubating overnight (approximately 20 h), followed by assays that measured their final optical density at 600 nm (OD_600_), resting pH, CFU/ml counts by serial dilution and plating, extracellular DNA (eDNA) levels by mixing with a DNA-specific fluorescent dye (SYTOX Green, Invitrogen)(91), and extracellular organic acids. Intracellular polysaccharide (IPS) compounds in the form of glycogen were measured by harvesting bacterial cells from exponential or stationary phase, followed by a colorimetric assay that combined hydrolysis of IPS in an alkaline solution with subsequent detection by the iodine-glycogen reaction (92).

### Plate-based competition assays

Overnight bacterial cultures were prepared using BHI medium in ambient atmosphere maintained with 5% CO_2_, diluted into TY medium containing 20 mM of glucose, galactose, GlcNAc, or GlcN, 10 mM each of glucose and galactose, or 10 mM of lactose, and incubated till OD_600_ reached 0.5. Six µl of *S. gordonii* culture was placed on a TY agar plate containing the same carbohydrate(s), and incubated for 24 h, followed by *S. mutans* which was similarly prepared and placed to the right of the colony. The plates were incubated for another day before photographing.

### Mixed-species biofilm assays

The role of ManLMNO of *S. sanguinis* SK36 in the interactions with *S. mutans* UA159 was assessed in an *in vitro,* mixed-species biofilm model. *S. sanguinis* was inoculated first. Briefly, SK36 and SK36/*manL* (n = 4, each with an Em-resistance marker) were grown overnight in BHI and sub-cultured by diluting 20-fold into fresh BHI, incubated at 37°C in an ambient atmosphere maintained with 5% CO_2_. Once OD_600_ reached 0.5, the cultures were diluted 100-fold into a BM medium (pH 7.0) (86). For standard condition, the BM medium contained 72 mM potassium phosphate buffer, 2 mM sucrose, and 18 mM glucose (BMGS). To study the effects of environmental factors, BMGS was modified to contain 5 mM arginine-HCl, 1 mM or 5 mM sodium pyruvate, reduced phosphate buffer (from 72 mM to 10 mM), or more glucose (from 18 mM to 1% or 55 mM). The cell suspensions were then aliquoted, at 400 µl/well, into individual wells of an ibidi^®^ µ-Slide 8-well chamber slide (#80827, ibidi GmbH). The chambers were pretreated with filter-sterilized human saliva (at 80 µl/well) at 37°C for 1 h, at which point the extra liquid was removed by aspiration. The cultures were incubated at 37°C in a 5% CO_2_ aerobic atmosphere, or an anaerobic jar (AnaeroPack containing the BD GasPack EZ system, resulting in an atmosphere containing <1% O_2_ and >13% CO_2_) for 24 h, refreshed with BMGS or any specified version of BMGS, and inoculated (4 µl/well) with the overnight culture of a *S. mutans* strain UA159-Km, which contains a Km-resistance marker in a non-essential gene (93), followed by another 2 days of incubation under the same conditions, with one more round of medium refreshment in between. At the end of the incubation, the culture supernatant was removed for measurement of pH values. The biofilm was washed 3 times with BM base medium and resuspended with 400 µl of the same medium. The biofilm was then scraped off by using a sterile pipette tip at medium force for at least 60 sec, transferred into a 1.7-ml centrifuge tube and sonicated for 15 sec at 100% power (FB120 water bath sonicator, Fisher Scientific), followed by serial dilution using PBS and plating on selective agar plates for CFU enumeration.

### RNA extraction, deep sequencing (RNA-seq), data analysis

5 m of bacterial cultures were prepared with TY medium containing 20 mM of glucose and harvested at mid-exponential phase (OD_600_ = 0.5∼0.6) and treated with RNAprotect reagent. Bacterial cells were resuspended in a lysis buffer (50 mM Tris-HCl pH 8.0, 10 mM EDTA, and 0.4% SDS) and disrupted by vigorous shaking for 1 min together with an equal volume of acidic phenol: chloroform (1:1, v/v) and similar amount of glass beads. After 10 min of centrifugation at 15,000× *g* at room temperature, the clarified aqueous layer was removed and processed using an RNeasy mini kit (Qiagen, Germantown, MD) for extraction of total RNA. Genomic DNA contamination was removed in column using an RNase-free DNase I solution (Qiagen). RNA deep sequencing for SK36 and the *manL* mutant was carried out by Microbial Genome Sequencing Center (MiGS, Pittsburgh, PA). The analysis of the RNA-Seq data was conducted on package R (94), following an established protocol detailed elsewhere (95). The compositional matrix of the expression data was normalized with the “voom” (96) function from the R package “limma” (97). The statistical analysis of the expression data was conducted by using the DESeq2 pipeline. A false discovery rate (FDR) of 0.01 and a fold-of-change of 2.0 were used as the cutoff values for identification of genes with differential expression. Findings from this analysis were highly consistent with RT-qPCR results included in our previous publication (32) and a few further confirmations were carried out but not shown here.

### Targeted metabolomics and biochemical assays for measurement of metabolites

Overnight cultures of strains SK36 and *manL* in BHI were diluted 20-fold into TY containing 20 mM glucose, and sub-cultured till OD_600_ reached 0.5. Immediately after centrifugation at 4°C at 10,000× *g* for 5 min, cell pellets were frozen at −80°C till analysis. Concentrations of 8 organic acids in bacterial extract, namely lactate, pyruvate, succinate, 3-hydroxybutyrate, α-ketoglutarate, malate, citrate, and fumarate were measured using liquid chromatography coupled with tandem mass spectrometry (LC-MS/MS) and internal controls. The results were normalized against the try weight of the samples.

The concentrations of pyruvate in bacterial culture supernatant were measured using a lactate dehydrogenase (Ldh)-based assay that coupled reduction of pyruvate with the oxidation of NADH, which was monitored as changes in optical density at a wavelength of 340 nm (OD_340_) using a UV spectrophotometer (32). A colorimetric method was used to measure acetoin in cultures (98). Briefly, 10 µl of bacterial supernate was mixed with 125 µl of a color reagent that was prepared by mixing equal volumes of 0.2% creatine solution in water and 1% α-naphthol freshly dissolved in 2.5 M sodium hydroxide. The reaction was incubated at room temperature for 40 min before the optical density (OD_525_) was measured. The concentrations of lactate, acetate, formate, and citrate were measured using a lactate assay kit (LS-K234, LSBio, Seattle, WA), an acetate assay kit (MAK086, Sigma), a formate assay kit (EFOR-100, BioAssay systems, Hayward, CA), and a citrate assay kit (MAK057, Sigma), respectively, by following instructions provided by the manufacturers. To avoid contaminating metabolites from yeast extract, most of these cultures were prepared using TV-glucose medium, although similar results were obtained for assays performed using both TV-glucose and TY-glucose. Each biochemical assay was conducted using at least three biological replicates, alongside a standard prepared using known concentrations of the substrate of interest.

### Statistics

Statistical analysis of data was carried out using the software of Prism (GraphPad of Dotmatics, San Diego, CA).

## Data Availability

The high-throughput sequencing data from this study have been deposited with Gene Expression Omnibus (GEO) and assigned accession number GSE209672.

## Acknowledgements

This study was supported by a grant from NIDCR to Lin Zeng and Robert A. Burne (DE012236). Metabolomic analysis of organic acids in bacteria was carried out by the metabolomics core at the Translational Research Institute at Advent Health in Orlando, Florida.

## Author Contributions

LZ designed the study; LZ and ZAT performed the experiments; ARW and LZ analyzed the data; LZ and RAB wrote the manuscript.

